# Radiographic and clinical risk factors of total knee arthroplasty in asymptomatic knees; a 15-year follow-up study

**DOI:** 10.1101/816462

**Authors:** Flemming Kromann Nielsen, Anne Grethe Jurik, Anette Jørgensen, Niels Egund

## Abstract

**Background:** Radiographic signs of osteoarthritis (OA) are frequent in knees without symptoms. The long-term impact of these findings is not completely elucidated. We wanted to evaluate whether radiographic or clinical baseline findings are associated with the risk of total knee arthroplasty (TKA) in knees without symptomatic OA but with clinical OA of the other knee during a mean follow-up period of 15 years.

**Methods:** A follow-up analysis was performed in 100 persons with unilateral, clinical knee OA according to the ACR (American College of Rheumatology) criteria, who participated in a clinical trial between 2000 and 2002. Baseline radiographs of the contralateral, non-symptomatic knee were available in 88 participants at follow-up. Data on TKA procedures were extracted from the Danish National Patient Register at follow-up. Radiographic and clinical findings were analyzed for associations with subsequent TKA.

**Results:** At follow-up, 40% had received a TKA in their non-symptomatic knee. The risk of TKA was significantly associated with baseline joint space narrowing (risk ratio (RR) 1.6 (95% confidence interval (95% CI) 1.4 to 1.9)), osteophytes (RR 1.5 (95% CI 1.3 to 1.8)) and subchondral sclerosis (RR 2.4 (95% CI 1.6 to 3.7)). Among the clinical findings, only baseline body mass index (BMI) was significantly associated with the risk of TKA (RR 1.4 (95& CI 1.1 to 1.8)).

**Conclusions:** Radiographic OA changes and BMI at baseline were significantly associated with the long-term risk of TKA in persons without symptomatic knee OA but with symptomatic OA in the contralateral knee, implying that radiographic OA findings are important prognostic factors regardless of symptoms.

## Introduction

Osteoarthritis (OA) of the knee is one of the most common causes of debilitating joint disease (1). Moreover, the economic burden associated with treatment and loss of productivity is substantial (2). The course of disease varies considerably, and it is essential to identify valid biomarkers that can be used to accurately predict disease progression. Previous studies have described potential biochemical markers (3,4) but imaging has the advantage of visualizing OA in specific joints.

Radiography and magnetic resonance imaging (MRI) are the most widely used imaging techniques in knee OA; the latter primarily for scientific purposes. Thus, MRI has gathered considerable attention in knee OA research, partly due to its ability to detect pre-clinical OA. However, there is currently no formalized MRI-based OA definition (5,6).

Radiography is an established method in the diagnosis of knee OA (7) and is often used in disease follow-up (8). An inconsistent association between radiographic and symptomatic OA, however, hampers diagnosis and complicates management of patients with knee OA (9,10). Thus, clinical signs of OA can be seen without radiographic OA and radiographic OA changes are frequently observed in persons without symptoms (11). However, increasing radiographic OA severity is more likely to be accompanied by symptoms (12).

The long-term consequences of radiographic changes in symptomatic knee OA have been investigated in several studies (13–15) whereas the significance of such findings in asymptomatic knees remains uncertain. Radiographic OA might, however, be an important prognostic marker in asymptomatic knees and could be associated with the long-term risk of symptomatic OA development and eventually total knee arthroplasty (TKA).

In a previous study, radiographic changes, specifically joint space narrowing (JSN), were associated with the risk of disease progression and TKA in patients with symptomatic knee OA during a follow-up period of 15 years (15). The contralateral, non-symptomatic knee was examined by radiography at baseline, but not investigated further. Follow-up data on TKA was obtained for both knees (15).

In this study, we investigated whether radiographic or clinical findings at baseline are associated with the incidence of TKA in knees without symptomatic OA during a mean follow-up period of 15 years in persons with symptomatic OA of the other knee.

## Materials and methods

### Data sources

The study was partly based on data from the Danish Civil Registration System and the Danish National Patient Register.

The Danish Civil Registration System assigns a unique 10-digit personal identification number to all Danish citizens, enabling accurate linkage across all national registers. The Danish National Patient Register includes data on all hospital in-patient as well as out-patient contacts. Danish public and private hospitals are obliged to report discharge diagnoses to the Danish National Patient Register, which are coded according to the Danish version of the International Classification of Diseases (ICD-10).

### Participants

Participants were recruited from a previous multi-center, randomized, placebo-controlled trial comparing the effect of five intra-articular injections of Hyalgan® and placebo (16) in 337 participants fulfilling the clinical criteria for knee OA according to the ACR (American College of Rheumatology) criteria (7). The inclusion criteria were primary knee OA, a Lequesne Algofunctional Index score of 10 or more (range: 0-24), a normal C-reactive protein level and e-GFR ≥ 60. Exclusion criteria were symptomatic OA from the other knee, inflammatory joint disease or severe co-morbidity (such as cancer and poor general health). Radiographic OA changes of the non-symptomatic was not an exclusion criterion. The participants entered the study between January 2000 and December 2002.

A total of 102 participants entered the study at our institution and radiography was obtained of both knees. At follow-up, radiographs were missing in 14 non-symptomatic knees and in two target knees; thus, evaluation of radiographic changes in the non-symptomatic knee was confined to 88 participants. The 100 target knees have already been investigated in a previous study (15).

Demographic variables were measured at baseline. Participants filled out a Western Ontario and McMaster Universities (WOMAC) questionnaire (17) regarding their symptomatic knee. The WOMAC score is divided in three domains: Pain (five items), stiffness (two items) and physical function (17 items). We used a 100 mm Visual Analogue Scale ranging from no pain/stiffness/difficulty (0) to extreme pain/stiffness/difficulty (100) with the following ranges: pain = 0-500, stiffness = 0-200, physical function = 0-1700, total score = 0-2400. No symptomatic scoring was performed of the 88 non-symptomatic knees.

The original study was approved by the Central Denmark Region Committee on Health Research Ethics and was carried out in accordance with the Declaration of Helsinki. Written informed consent was obtained from all participants prior to participation, including consent to publish study results and images.

The Central Denmark Region Committee on Health Research Ethics was contacted prior to this follow-up study and the need for ethical approval was waived. The study was approved by the Danish Data Protection Agency (1-16-02-126-16).

### Imaging and image analysis

Radiographs of both knees were obtained at study inclusion using a one leg weight-bearing technique in 30^0^ flexion in three views: Posteroanterior and lateral views of the tibiofemoral, and axial view of the patellofemoral joint space. The initial lateral radiograph was used as guidance for securing optimal inclination of the lower leg for visualization of the tibiofemoral and patellofemoral joint spaces (18).

All radiographs were anonymized prior to analysis and the reader was blinded to study information and outcome. Radiographic evaluations were performed by a senior musculoskeletal radiologists (NE) according to a modified Ahlbäck grading (19); Grade 0 = normal joint spaces, grade 0.5 = minimal but definite JSN (25% JSN), grade 1.0 = > 50%, grade 1.5 = > 75% and grade 2.0 = 100% JSN; grade 3 = 100% JSN and < 5 mm bone attrition; grade 4 = 100% JSN and > 5 mm bone attrition. We defined OA as JSN ≥ grade 0.5 in at least one knee joint space. Osteophytes (grades 0-3), subchondral sclerosis (0/1) and subchondral cysts (0/1) were also assessed (20). Osteophytes were graded as follows: grade 0 = no osteophytes, grade 1 = possible or small osteophytes, grade 2 = definite and moderate osteophytes, grade 4 = large osteophytes.

### Acquisition of follow-up data

Data on TKA procedures in both knees were extracted from the Danish National Patient Register, dividing the study population into a TKA and a non-TKA group.

### Statistical analysis

Data were analyzed using Stata 13.0 (StataCorp, College Station, TX, USA). Baseline characteristics were analyzed descriptively using means and standard deviations (SDs).

Our primary goal was to analyze any associations between baseline radiographic changes and the risk of TKA in the non-symptomatic knee at follow-up. Furthermore, to analyze if other baseline variables, including sex, age and body mass index (BMI), as well as the degree of symptoms in the inclusion knee, were associated with the risk of TKA in the non-symptomatic knee.

Associations were analyzed using univariable and multivariable log-binominal regression analyses. Multivariable log-binominal regression was performed for radiographic measures only, with adjustment for possible confounders (sex, age, BMI). The extent of any significant association was expressed in risk ratios (RR) with 95% confidence intervals (95% CI).

Kaplan-Meier plots were made comparing the cumulative incidence of TKA over time in relation to JSN (21). BMI was divided into four grades for calculations of RR: Grade 1 (normal) = <25 kg/m^2^, Grade 2 (overweight) = 25 -<30 kg/m^2^, Grade 3 (obese) = 30 -<35 kg/m^2^ and Grade 4 (very obese) = ≥35 kg/m^2^.

## Results

Baseline characteristics of the 88 participants are shown in Table 1. The mean follow-up time was 15.2 years (range 14.2 to 16.4 years). A total of 35 participants had undergone TKA in the non-symptomatic knee at follow-up. Among the different baseline demographic and clinical variables, only an increased BMI was significantly associated with the risk of subsequent TKA, RR 1.4 (95% CI 1.1 to 1.8).

**Table 1.** Baseline characteristics of the 88 participants and total knee arthroplasty of the non-symptomatic knee.

| Variable | Total (n=88) | Total knee arthroplasty (TKA)* |  | p Value | If p value <0.05<br>RR (95% CI) |
| --- | --- | --- | --- | --- | --- |
|  |  | Yes (n=35) | No (n=53) |  |  |
| Female, % (no.) | 69.3% (61) | 65.7% (23) | 71.7% (38) | 0.53 | - |
| Age (years), mean (SD) | 61.6 (9.9) | 59.6 (9.3) | 62.3 (10.2) | 0.15 | - |
| BMI (kg/m<sup>2</sup>), mean (SD) | 28.7 (4.6) | 30.3 (4.3) | 27.7 (4.5) | 0.003 | 1.4 (1.1 to 1.8) |
| WOMAC, mean (SD): |  |  |  |  |  |
| - *Pain* | 182 (90) | 184 (96) | 181 (87) | 0.87 | - |
| - *Stiffness* | 80 (44) | 89 (44) | 74 (43) | 0.08 | - |
| - *Function* | 656 (318) | 684 (337) | 637 (307) | 0.48 | - |
| Hyalgan treatment %, (no.)** | 46.6% (41) | 45.7% (16) | 47.2% (25) | 0.89 | - |

RR: risk ratio; CI: confidence interval; RR: relative risk; SD: standard deviation; BMI: body mass index; WOMAC: Western Ontario and McMaster Universities OA index
\* Number of participants with TKA at follow-up in knees without symptomatic knee osteoarthritis at baseline
\*\*Hyalgan treatment was administered locally in the symptomatic knee (i.e. the inclusion knee)

The radiographic findings at baseline in the 88 non-symptomatic knees are shown in Table 2. Radiographic OA changes were frequent: 66% had at least Grade 0.5 JSN, osteophytes of any grade were present in 52%, while subchondral sclerosis was present in 13%. Subchondral cysts were only present in three knees (one with subsequent TKA) and were omitted from further analysis due to the low prevalence.

**Table 2.**
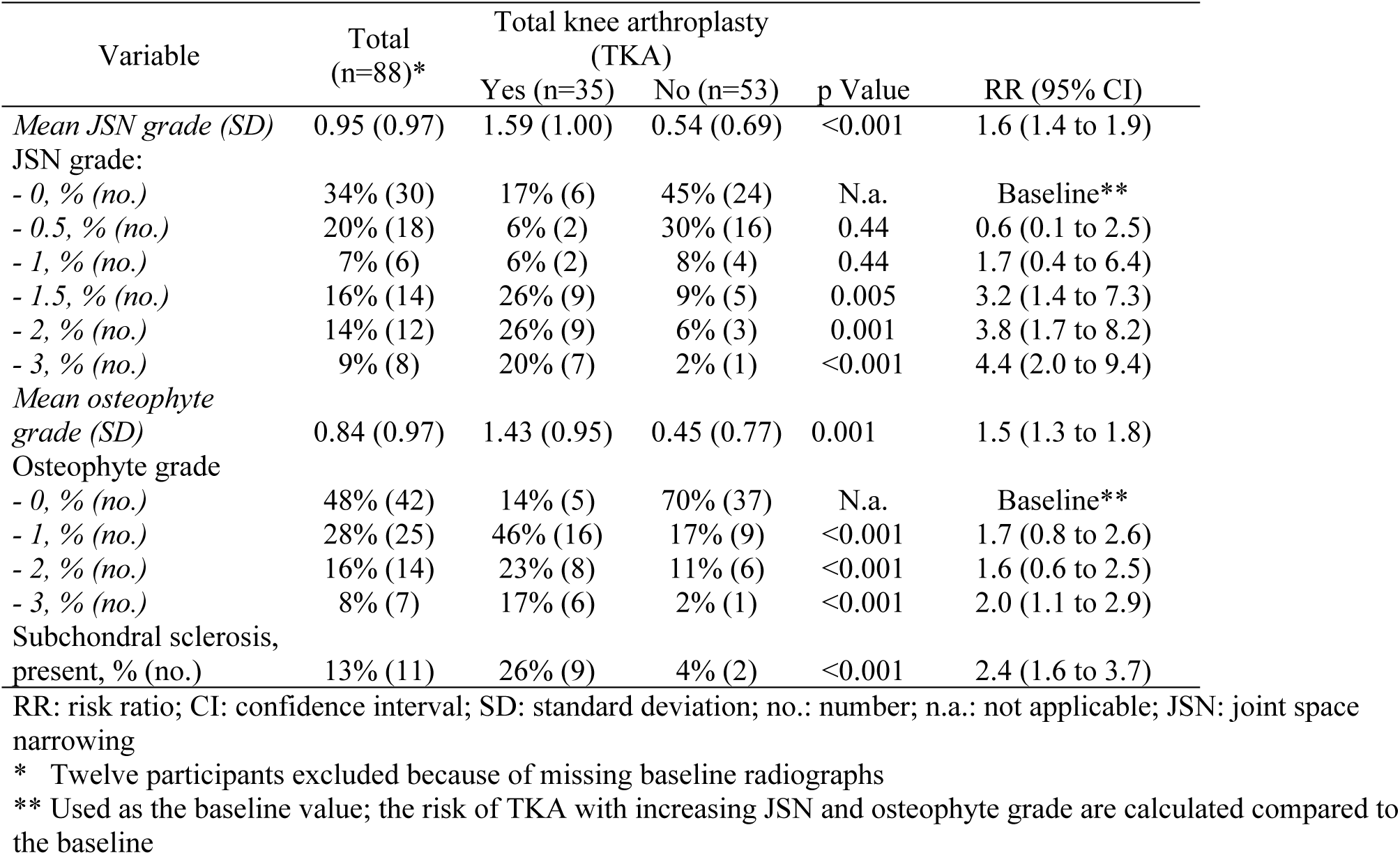
Radiographic predictors for knee replacement in knees without symptomatic knee OA at baseline

| Variable | Total<br>(n=88)* | Total knee arthroplasty<br>(TKA) |  | p Value | RR (95% CI) |
| --- | --- | --- | --- | --- | --- |
|  |  | Yes (n=35) | No (n=53) |  |  |
| *Mean JSN grade (SD)* | 0.95 (0.97) | 1.59 (1.00) | 0.54 (0.69) | <0.001 | 1.6 (1.4 to 1.9) |
| JSN grade: |  |  |  |  |  |
| - 0, % (no.) | 34% (30) | 17% (6) | 45% (24) | N.a. | Baseline** |
| - 0.5, % (no.) | 20% (18) | 6% (2) | 30% (16) | 0.44 | 0.6 (0.1 to 2.5) |
| - 1, % (no.) | 7% (6) | 6% (2) | 8% (4) | 0.44 | 1.7 (0.4 to 6.4) |
| - 1.5, % (no.) | 16% (14) | 26% (9) | 9% (5) | 0.005 | 3.2 (1.4 to 7.3) |
| - 2, % (no.) | 14% (12) | 26% (9) | 6% (3) | 0.001 | 3.8 (1.7 to 8.2) |
| - 3, % (no.) | 9% (8) | 20% (7) | 2% (1) | <0.001 | 4.4 (2.0 to 9.4) |
| *Mean osteophyte grade (SD)* | 0.84 (0.97) | 1.43 (0.95) | 0.45 (0.77) | 0.001 | 1.5 (1.3 to 1.8) |
| Osteophyte grade |  |  |  |  |  |
| - 0, % (no.) | 48% (42) | 14% (5) | 70% (37) | N.a. | Baseline** |
| - 1, % (no.) | 28% (25) | 46% (16) | 17% (9) | <0.001 | 1.7 (0.8 to 2.6) |
| - 2, % (no.) | 16% (14) | 23% (8) | 11% (6) | <0.001 | 1.6 (0.6 to 2.5) |
| - 3, % (no.) | 8% (7) | 17% (6) | 2% (1) | <0.001 | 2.0 (1.1 to 2.9) |
| Subchondral sclerosis,<br>present, % (no.) | 13% (11) | 26% (9) | 4% (2) | <0.001 | 2.4 (1.6 to 3.7) |

RR: risk ratio; CI: confidence interval; SD: standard deviation; no.: number; n.a.: not applicable; JSN: joint space narrowing
\* Twelve participants excluded because of missing baseline radiographs
\*\* Used as the baseline value; the risk of TKA with increasing JSN and osteophyte grade are calculated compared to the baseline

A significant association was found between TKA at follow-up and baseline JSN (RR 1.6 (95% CI 1.4 to 1.9)), osteophytes (RR 1.5 (95% CI 1.3 to 1.8)) and subchondral sclerosis (RR 2.4 (95% CI 1.6 to 3.7)). The associations did not change after adjustment for potential confounders (results not shown).

Increasing grades of JSN and osteophytes at baseline increased the risk of TKA at follow-up, see Figure 1 and Table 2. Thus, the RR of undergoing TKA in persons with grades 2 and 3 JSN was 3.8 (95% CI 1.7 to 8.2) and 4.4 (95% CI 2.0 to 9.4), respectively, compared to persons with JSN grade 0 (Table 2). The wide confidence intervals reflect the relatively low number of persons in each group. The Kaplan-Meier time to event analysis illustrates the relation between increasing degrees of JSN at baseline and the risk of TKA (Figure 1).

**Figure 1.**
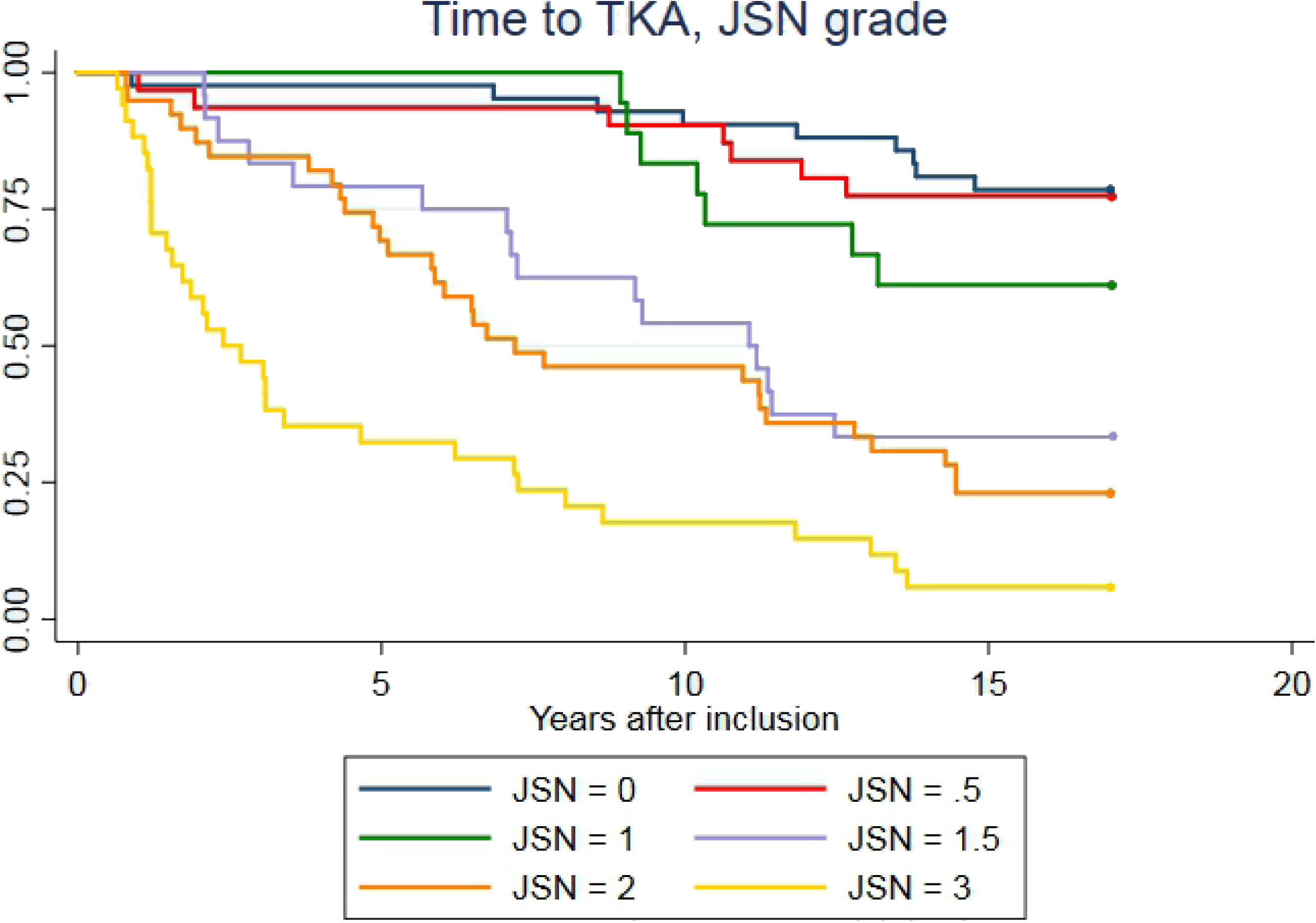
Kaplan-Meier plot showing the cumulative incidence of TKA during follow-up in the non-symptomatic knee in relation to different degrees of JSN at baseline TKA: total knee arthroplasty; JSN: joint space narrowing

Sixty-six participants had undergone TKA in their symptomatic knee at follow-up (results published elsewhere) (15). All participants who undergone a TKA in the non-symptomatic knee already underwent TKA in the symptomatic knee.

## Discussion

In this study we found that radiographic OA changes and BMI at baseline were associated with the long-term risk of TKA in non-symptomatic knees during a mean follow-up period of 15 years. Thus, radiographic OA should not be interpreted as a random finding in an otherwise asymptomatic individual but as a risk factor for future OA development, at least in persons with contralateral knee OA.

The prevalence of radiographic OA changes, frequent knee pain and symptomatic knee OA was recently examined in a random sample of 10,000 people between 56 and 84 years of age (mean age 70 years) (11). Frequent knee pain was found in 25.1%. A random sample of 1,300 individuals with frequent knee pain and 650 without were further analyzed. Radiographic OA (defined as Kellgren-Lawrence (K-L) grade of ≥ 2) was found in 25.6% and symptomatic knee OA (defined either as clinical knee OA according to the ACR clinical criteria or as radiographic knee OA and frequent knee pain) was found in 15.4%. Radiographic knee OA without symptoms were seen in 11% and even though these persons did not have OA according to the ACR criteria, our results indicate an increased long-term risk of TKA.

The natural course of persons without frequent knee pain but with radiographic knee OA has only been examined in a limited number of studies. In the US Framingham Osteoarthritis Study (22), 869 persons were followed for a mean period of 8.1 years. A total of 30% (n=261) had asymptomatic radiographic knee OA at baseline (KL ≥ 2) of which 29% (n=76) had radiographic progression at follow-up; 11% (n=93) developed asymptomatic radiographic knee OA during follow-up. The number of TKAs was not reported. In the UK Chingford Women’s Study (23), 561 asymptomatic women were followed for a mean period of 14 years. A total of 19% (n=106) had radiographic knee OA at baseline (KL ≥ 2) of which 38% (n=40) had radiographic progression at follow-up. Nineteen had received a TKA at follow-up; six of them had a baseline K-L ≥2 and 13 had a baseline K-L <2.

Our study sample differed from the populations in these community-based studies (22,23) as symptomatic knee OA was already present in the contralateral knee. Furthermore, our definition of radiographic knee OA was based on the degree of joint space narrowing (19,24) whereas the K-L grading was based on the presence of osteophytes (25). Thus, K-L grade 2 used to define radiographic knee OA in these studies (22,23) was defined as a “definite osteophyte, possible JSN” (23).

Despite these differences, our results were similar. Radiographic knee OA was seen in 66% (n=58) at baseline (defined as any degree of JSN); 40% (n=35) progressed to TKA of which 33% (n=29) had some degree of JSN at baseline and 7% (n=6) did not (Table 2). Thus, the rate of disease progression was similar to our results, indicating that the risk of symptomatic knee OA is significant once radiographic signs of OA are present. Furthermore, our results showed that the severity of radiographic changes highly increased the risk of TKA and shortened the time before TKA was performed, see Figure 1 and Table 2.

Many risks factors for knee OA are considered systemic, including age, sex and BMI (26), and thus affect both knees. However, in a study by McMahon and Block examining the rate of contralateral knee replacement after TKA, no association was found between any systemic risk factor, including age, sex and BMI, and the subsequent risk of TKA (27). Instead, the baseline K-L grade was strongly correlated with the future risk of TKA, supporting our main finding, namely that the degree of radiographic OA changes is an important determinant for the future risk of TKA.

We did however also find an association between BMI and the risk of TKA in the non-symptomatic knee. It has been hypothesized that increased BMI might be more important in the development of symptomatic knee OA than in disease progression once symptomatic knee OA is present (28). Our findings seem to corroborate this hypothesis. Thus, a previous analysis of our study population revealed that an increased BMI was not associated with the risk of TKA in the symptomatic knee (15). The present analysis of the non-symptomatic knee shows an association between BMI and the risk of developing symptomatic knee OA resulting in TKA. The study by McMahon and Block (27) did not report whether the contralateral knee was asymptomatic at baseline and symptoms might thus already have been present. This would minimize the importance of an increased BMI and help explain the differing results.

Interestingly, all 35 participants who underwent TKA in the non-symptomatic knee had already undergone TKA in the symptomatic knee. This concurrence may partly be explained by the effect of TKA: persons who have a TKA in one knee due to symptomatic knee OA find that this treatment is helpful and are more likely to accept a TKA in the contralateral knee if symptoms of knee OA should occur.

Our study has some limitations. The number of participants was limited compared to other longitudinal studies (29). Only radiographs were available of the non-symptomatic knee. MR examinations would undoubtedly have yielded additional information and might have shown other significant associations as demonstrated previously (15). Our analysis was limited to baseline radiographic data, and no clinical or radiological follow-up data were available including BMI. Thus, we did not know whether any of these parameters changed during the follow-up period. Using TKA as a clinical endpoint can be problematic since the final decision to perform a TKA is based on many individual factors including local surgical practices and patient preferences. The radiographs were only assessed by one reader. However, high observer agreement was found in a previous study (30). Among the strengths of our study is the use of highly reliable register data on TKA combined with a long follow-up period.

## Conclusions

We found that radiographic OA changes as well as increased BMI were important prognostic markers for the future risk of TKA in persons without symptomatic knee OA but with symptomatic OA in the contralateral knee, implying that radiographic OA findings are important prognostic factors regardless of symptoms.

## Acknowledgements

We thank Kristian Stengaard-Pedersen for collecting patient material.

## References

1. Cross M, Smith E, Hoy D, Nolte S, Ackerman I, Fransen M, et al. The global burden of hip and knee osteoarthritis: estimates from the global burden of disease 2010 study. Ann Rheum Dis. 2014 Jul;73:1323–30.

2. Salmon JH, Rat AC, Sellam J, Michel M, Eschard JP, Guillemin F, et al. Economic impact of lower-limb osteoarthritis worldwide: a systematic review of cost-of-illness studies. Osteoarthr Cartil. 2016;24:1500–8

3. Williams FM, Spector TD. Biomarkers in osteoarthritis. Arthritis Res Ther. 2008;10:101.

4. M Lotz et al. Value of biomarkers in osteoarthritis: current status and persepctives. Ann Rheum Dis. 2013 Nov;72:1756–63.

5. Hunter DJ, Arden N, Conaghan PG, Eckstein F, Gold G, Grainger A, et al. Definition of osteoarthritis on MRI: results of a Delphi exercise. Osteoarthr Cartil. 2011 Aug;19:963–9.

6. Menashe L, Hirko K, Losina E, Kloppenburg M, Zhang W, Li L, et al. The diagnostic performance of MRI in osteoarthritis: a systematic review and meta-analysis. Osteoarthr Cartil. 2012 Jan;20:13–21.

7. Altman R, Asch E, Bloch D, Bole G, Borenstein D, Brandt K, et al. Development of criteria for the classification and reporting of osteoarthritis. Classification of osteoarthritis of the knee. Diagnostic and Therapeutic Criteria Committee of the American Rheumatism Association. Arthritis Rheum. 1986 Aug;29:1039–49.

8. Hunter DJ, Felson DT. Osteoarthritis. BMJ. 2006 Mar;332:639–42.

9. Dieppe PA, Cushnaghan J, Shepstone L. The Bristol “OA500” study: progression of osteoarthritis (OA) over 3 years and the relationship between clinical and radiographic changes at the knee joint. Osteoarthr Cartil. 1997 Mar;5:87–97.

10. Hochberg MC, Lawrence RC, Everett DF, Cornoni-Huntley J. Epidemiologic associations of pain in osteoarthritis of the knee: data from the National Health and Nutrition Examination Survey and the National Health and Nutrition Examination-I Epidemiologic Follow-up Survey. Semin Arthritis Rheum. 1989 May;18:4–9.

11. Turkiewicz A, Gerhardsson M, Engstrom G, Nilsson PM, Mellstrom C, Lohmander LS, et al. Prevalence of knee pain and knee OA in southern Sweden and the proportion that seeks medical care. Rheumatology (Oxford). 2015;54:827–35.

12. Bedson J, Croft PR. The discordance between clinical and radiographic knee osteoarthritis: a systematic search and summary of the literature. BMC Musculoskelet Disord. 2008 Sep;9:116.

13. Chapple CM, Nicholson H, Baxter GD, Abbott JH. Patient characteristics that predict progression of knee osteoarthritis: a systematic review of prognostic studies. Arthritis Care Res. 2011 Aug;63:1115–25.

14. Riddle DL, Kong X, Jiranek WA. Factors associated with rapid progression to knee arthroplasty: complete analysis of three-year data from the osteoarthritis initiative. Joint Bone Spine. 2012;79:298–303.

15. Nielsen FK, Egund N, Jørgensen A, Jurik AG. Risk factors for joint replacement in knee osteoarthritis; a 15-year follow-up study. BMC Musculoskelet Disord. 2017 Dec;18:510.

16. Jorgensen A, Stengaard-Pedersen K, Simonsen O, Pfeiffer-Jensen M, Eriksen C, Bliddal H, et al. Intra-articular hyaluronan is without clinical effect in knee osteoarthritis: a multicentre, randomised, placebo-controlled, double-blind study of 337 patients followed for 1 year. Ann Rheum Dis. 2010 Jun;69:1097–102.

17. Bellamy N. Western Ontario and McMaster Universities Osteoarthritis Index (WOMAC). http://www.rheumatology.org/I-Am-A/Rheumatologist/Research/Clinician-Researchers/Western-Ontario-McMaster-Universities-Osteoarthritis-Index-WOMAC. Accessed 08/17 2017.

18. Skou N, Egund N. Patellar position in weight-bearing radiographs compared with non-weight-bearing: significance for the detection of osteoarthritis. Acta Radiol. 2017 Mar;58:331–7.

19. Ahlback S. Osteoarthrosis of the knee. A radiographic investigation. Acta Radiol Diagn (Stockh). 1968;Suppl 277:7–72.

20. Egund N, Ryd L. Patellar and quadriceps mechanism. In: Davies AM C-P V, editor. Berlin, Heidelberg: Springer; 2002. p. 217–48. (Imaging of the Knee: Techniques and Applications.).

21. Kaplan EL, Meier P. Nonparametric Estimation from Incomplete Observations. J Am Stat Assoc. 1958;53:457–81.

22. Felson DT, Zhang Y, Hannan MT, Naimark A, Weissman BN, Aliabadi P, et al. The incidence and natural history of knee osteoarthritis in the elderly, the framingham osteoarthritis study. Arthritis Rheum. 1995 Oct;38:1500–5.

23. Leyland KM, Hart DJ, Javaid MK, Judge A, Kiran A, Soni A, et al. The natural history of radiographic knee osteoarthritis: a fourteen-year population-based cohort study. Arthritis Rheum. 2012 Jul;64:2243–51.

24. Boegard T, Rudling O, Petersson IF, Sanfridsson J, Saxne T, Svensson B, et al. Postero-anterior radiogram of the knee in weight-bearing and semiflexion. Comparison with MR imaging. Acta Radiol. 1997 Nov;38:1063–70.

25. Kellgren JH, Lawrence JC. Radiological assessment of osteo-arthrosis. Ann Rheum Dis. 1957 Dec;16:494–502.

26. Felson DT, Zhang Y, Hannan MT, Naimark A, Weissman B, Aliabadi P, et al. Risk factors for incident radiographic knee osteoarthritis in the elderly: the Framingham Study. Arthritis Rheum. 1997 Apr;40:728–33.

27. McMahon M, Block JA. The risk of contralateral total knee arthroplasty after knee replacement for osteoarthritis. J Rheumatol. 2003 Aug;30:1822–4.

28. Cooper C, Snow S, McAlindon TE, Kellingray S, Stuart B, Coggon D, et al. Risk factors for the incidence and progression of radiographic knee osteoarthritis. Arthritis Rheum. 2000;43:995–1000.

29. Roemer FW, Kwoh CK, Hannon MJ, Hunter DJ, Eckstein F, Wang Z, et al. Can structural joint damage measured with MR imaging be used to predict knee replacement in the following year? Radiology. 2015 Mar;274:810–20.

30. Rytter S, Egund N, Jensen LK, Bonde JP. Occupational kneeling and radiographic tibiofemoral and patellofemoral osteoarthritis. J Occup Med Toxicol. 2009 Jul;4:19.

